# Early-life telomere length variation under changing developmental conditions in long-lived bats

**DOI:** 10.64898/2026.05.26.727814

**Authors:** Tadhg Lonergan, Megan L. Power, Luke Romaine, Roger D. Ransome, Frédéric Touzalin, Sebastien J. Puechmaille, Gareth Jones, Emma C. Teeling

**Affiliations:** School of Biology and Environmental Science, Science Centre East, University College Dublin, Belfield, Dublin 4, Ireland; School of Biological Sciences, University of Bristol, Life Sciences Building, 24 Tyndall Avenue, Bristol BS8 1TQ, United Kingdom; School of Biodiversity, One Health and Veterinary Medicine, University of Glasgow, Glasgow, United Kingdom; ISEM, University of Montpellier, CNRS, IRD, Montpellier, France; Institut Universitaire de France, Paris, France

**Keywords:** Telomeres, Chiroptera, Early-life environment, Climate, Development, Maternal age, Weather conditions

## Abstract

Early-life conditions can shape molecular ageing processes, yet to what extent developmental variation in telomere length (TL) influences ageing trajectories remains unclear, particularly in long-lived mammals. We investigated how early-life environmental conditions and maternal age relate to juvenile TL and short-term survival in two long-lived bat species, *Myotis myotis* and *Rhinolophus ferrumequinum*. Using novel long-term datasets spanning ten years in *M. myotis* and five years in *R. ferrumequinum*, we measured relative telomere length (rTL) in juvenile wing tissue and applied sliding window analysis to identify sensitive climatic periods during development. In both species, early-life rTL varied significantly among years and was associated with short-term climatic conditions, with rainfall predicting rTL in both species and temperature acting in opposing directions: longer rTL with warmer conditions in *M. myotis*, and longer rTL at intermediate temperatures in *R. ferrumequinum*. Maternal age at conception showed little association with offspring TL in either species, although a weak positive sex-specific longitudinal effect was detected in *R. ferrumequinum*. Despite clear environmental influences on early-life rTL, we found no evidence that early-life rTL or early-life telomere change predicted short-term survival. Together, these results indicate that early-life telomere variation in bats reflects climatic conditions during development, providing novel insights into how early-life exposures could contribute to inter-individual differences in ageing trajectories in long-lived mammals.

## Introduction

Early-life conditions can have significant short-term effects on and long-lasting consequences for an individual’s fitness (1), often manifested as declines in reproductive success and/or survival late in the lifespan (2–4). While the mechanisms behind such patterns remain unclear, telomere length (TL) has been proposed as a biomarker to link early-life conditions with late-life declines in fitness parameters (5). Telomeres are conserved repetitive nucleotide sequences (TTAGGG) that cap the ends of linear chromosomes, preserving genomic stability during cell division (6). Due to the end-replication problem and exposure to oxidative damage, telomeres progressively shorten with cell division and physiological stress (7–9). When telomeres become critically short, cells activate a DNA damage response leading to replicative senescence or apoptosis (10), processes that contribute to tissue degeneration and organismal ageing (11,12). Telomere attrition is therefore recognised as one of the primary hallmarks of ageing (13), although whether telomere shortening acts predominantly as a causal driver of ageing or as a downstream consequence of accumulated cellular damage remains an active area of debate (14–16).

Telomere dynamics are especially sensitive during early development, a period of rapid growth and high cellular turnover during which substantial telomere shortening has been documented in many vertebrates (17–19) though a recent meta-analysis found that early-life attrition is not universally faster than adult attrition across taxa (20). Rapid early-life telomere attrition has been documented across diverse taxa including humans *Homo sapiens* (21), baboons *Papio spp.* (22), Soay sheep *Ovis aries* (23), European shags *Phalacrocorax aristotelis*, wandering albatrosses *Diomedea exulans* (24), and sand lizards *Lacerta agilis* (25). Moreover, early-life TL has been associated with survival and reproductive success in zebra finches *Taeniopygia guttata* (26) and Soay sheep (23), although patterns vary among systems, suggesting that both initial TL and the rate of early attrition may shape individual life-history trajectories (27,28).

Environmental conditions play a major role in shaping early-life telomere dynamics. Stressors such as nutritional limitation, extreme temperatures, and adverse weather can accelerate telomere shortening through increased metabolic demand and oxidative stress (29,30). Climatic variables, particularly temperature and rainfall, have emerged as consistent drivers of telomere variation. For example, TL in European badger (*Meles meles*) cubs is associated with spring weather conditions (28), and developmental temperature and precipitation influence telomere dynamics in sand lizards (*Lacerta agilis*) and several bird species, likely via effects on growth, provisioning, and physiological stress (25,31,32). As climate variability and extreme events intensify (33), it is important to understand how climatic conditions affect life-history strategies and how this is reflected in TL variation.

In addition to climatic effects, parental age at conception can shape offspring TL and early-life ageing trajectories. In humans and chimpanzees (*Pan troglodytes*), older fathers often produce offspring with longer telomeres, potentially due to persistent telomerase activity in the testes or selective survival of germ cells with longer telomeres (34–36). However, parental age effects are not universal: negative paternal age effects have been reported in several birds and mammals, while maternal age effects vary widely across taxa in both direction and magnitude (37–40). Parental age may influence offspring telomere dynamics through both germline mechanisms (e.g. older mothers producing higher-quality offspring due to selective oocyte usage (41)) and through age-related differences in parental condition and postnatal care, which could generate both beneficial and detrimental effects (42,43). Additionally, parental age effects on offspring may differ between offspring sexes (43,44). These complex and often inconsistent findings underscore the necessity for more extensive research, utilizing longitudinal datasets across diverse taxa, into the parental effects and the long-term fitness implications associated with parental age effects on early-life TL

Bats (Order Chiroptera) provide an exceptional model for investigating developmental influences on ageing. Despite their small body size and high metabolic rates, bats exhibit extreme longevity relative to other mammals of similar size with several species showing delayed or negligible senescence relative to predictions based on body size (45–48). Longevity in mammals is compared using the longevity quotient (LQ), defined as the ratio of observed maximum lifespan to that predicted for a mammal of equivalent body mass (45), which corrects for the confounding effect of body size. Bats consistently exhibit some of the highest LQs recorded among mammals, with many species living far longer than expected based on body size alone (49). Juvenile bats undergo rapid postnatal growth over a period of approximately 4 - 6 weeks, during which they remain dependent on maternal care and mother’s local foraging conditions, making early development a particularly sensitive window during which environmental and parental effects can have lasting physiological consequences (50–52). In temperate bat species, reproduction and juvenile development occur within narrow seasonal windows that impose strong constraints on growth and survival (53,54). Variation in spring and summer weather influences insect availability, maternal foraging success, juvenile growth trajectories and oxidative stress (29,51,55), thereby shaping energetic trade-offs between growth, maintenance, and cellular repair. These constraints are likely to magnify the impact of developmental environments on molecular ageing processes such as telomere dynamics.

Telomere dynamics have been investigated in a small number of bat species, with prior work predominantly focused on age-related telomere change in adults (56). In the long-lived *Myotis* genus, telomeres show no consistent attrition with chronological age, in both cross-sectional (*M. myotis*, *M. bechsteini*i, *M. lucifugus*; 57,58) and longitudinal analyses (*M. myotis*; 59). In contrast, telomeres shorten with age in *R. ferrumequinum* (57,60), although hibernation appears to slow or even reverse attrition, with within-individual telomere lengthening and increased telomerase expression observed across winter torpor (61). Beyond chronological age, environmental and life-history variables have also been linked to adult telomere variation in these populations: spring temperature, rainfall and wind speed predict longitudinal telomere change in adult *M. myotis* (59); climatic conditions during hibernation shape telomere dynamics in adult *R. ferrumequinum* (61); and reproductive timing and effort are associated with telomere length and survival in adult female *R. ferrumequinum* (60). Sex-specific patterns have also been reported in other bat species (62). However, to date the determinants of telomere variation during the narrow, physiologically demanding early-life developmental window have not been examined in bats.

Given that early-life conditions can have particularly pronounced and lasting effects on telomere dynamics in long-lived species, and juvenile bats face acute environmental constraints during their short growth window, understanding how early-life telomere variation is shaped by developmental conditions represents a distinct and important question for bat ageing biology. Here, we present the first detailed investigation of early-life telomere dynamics in bats, focusing on two long-lived temperate bat species: the greater mouse-eared bat (*Myotis myotis*) and the greater horseshoe bat (*Rhinolophus ferrumequinum*). *M. myotis* is among the longest-lived bats recorded, with individuals reaching at least 37 years of age in the wild (63) and exhibiting a high longevity quotient (LQ = 5.71). *R. ferrumequinum* exhibits similar exceptional longevity (maximum lifespan: 30 years old) and a comparable LQ (LQ = 4.98), with both species producing a single offspring annually and maintaining strong site fidelity across decades (54,64–66). Both species experience rapid juvenile growth and strong dependence on seasonal environmental conditions (51,55,67). Using long-term datasets spanning 10 years in *M. myotis* and 5 years in *R. ferrumequinum*, we test how early-life environmental conditions and parental age shape TL in juvenile bats. Specifically, we ask: (i) how TL changes within the early life developmental window, (ii) how climatic conditions during early life influence telomere dynamics, (iii) whether parental age predicts offspring TL, and (iv) whether juvenile TL predicts early survival. By integrating developmental, ecological, and parental influences on telomere biology, this study provides new insight into how early-life processes contribute to inter-individual differences in ageing trajectories in long-lived mammals.

## Materials and Methods

### (a) Study populations and sampling

Here we present a novel dataset of early-life TL measurements from two wild bat species; the greater mouse-eared bat (*Myotis myotis*) juveniles (∼8 weeks old: 500 individuals) sampled annually between 2014-2023 and 837 samples from 435 greater horseshoe bat individuals (*Rhinolophus ferrumequinum*; 0-78 days old) sampled across 2016-2020. These datasets target the early-life developmental period, a life-history stage not represented in previous bat telomere studies.

#### (i) Myotis myotis

Wing tissue was sampled from a wild population of the greater mouse-eared bat (*Myotis myotis*), located in Morbihan, Brittany, France. This population has been monitored since 2010 in collaboration with Bretagne Vivante, a grass roots conservation organization, with annual summer sampling taking place from 2013 onwards (48,57,59). In this study we focused on a single maternity colony that occupies the attic of a school building in Férel, Brittany (47◦28’ N, 2◦20’ W), to which females return every spring and summer. This colony was selected due to its consistent annual sampling and continuous automated monitoring using fixed antenna systems installed at the roost entrance and exit which record the identities of tagged individuals as they pass in and out of the roost. During each annual sampling period (2014-2023; 10 years), most individuals of the colony were captured and sampled as they emerged from the roost using customised harp traps. Following capture, bats were placed in individual cloth bags before processing. Newly captured individuals were fitted with Passive Integrated Transponders (PIT) tags (2.12x11mm, 0.1g, Trovan®), containing a unique 10-digit code which can be read using passive Radio-Frequency Identification (RFID) readers and facilitate the identification of individuals at each subsequent recapture.

Annual *M. myotis* sampling was carried out in early July when juveniles are approximately 8 weeks old. Sampling occurred only once per year, during which bats were aged, sexed and morphometric measurements were taken including forearm length (radius, to an accuracy of ± 0.1mm using Vernier callipers) and weight (to an accuracy of ± 0.1g using a precision weight balance). Juveniles were distinguished from adults based on size, weight and the degree of ossification of the epiphyseal cartilage in the metacarpal-phalangeal joints (67). In adults, these joints appear fused while in juveniles they appear unfused with a distinctive “gap”. Wing biopsies were taken using a sterile 3mm biopsy punch. Wing biopsy sampling is a standard nonlethal tissue sample protocol commonly used with bats for genetic analysis (68) with wing tissue shown to be a suitable proxy for global tissue telomere dynamics in bats (69). Samples were immediately flash frozen in liquid nitrogen in the field and later transferred to a -80°C freezer for long term storage. Fifty *M. myotis* juvenile wing punch samples were chosen randomly each year from 2014 to 2023 for a total of five hundred individuals.

#### (ii) Rhinolophus ferrumequinum

Wing tissue was sampled from a wild population of greater horseshoe bats (*Rhinolophus ferrumequinum*) located at Woodchester Mansion, Gloucestershire, UK (51°43′ N, 2°18′ W). This maternity roost has been ringed and monitored continuously since 1959 (54,64). Pups are born primarily in late June to mid-July, but births can occur as late as August (54). In this study, wing biopsy punches (3 mm) were collected from pups during primary summer catching sessions (late June–mid July) each year between 2016 and 2020 (5 years). Sampling sessions occurred every two to four days where pups were sexed and morphometric measurements taken including body mass (± 0.1 g using a precision balance) and forearm length (radius; ± 0.1 mm using Vernier callipers). Birth dates were estimated with high accuracy because pups were sometimes first encountered with fresh umbilical cords attached (i.e. born that day), enabling the construction of growth curves using forearm length data (64). Most pups have an initial wing biopsy taken within four days of birth. Additional sampling later within the same summer (August–September) was conducted for a subset of individuals to quantify within-individual changes in early life (Supplementary Table S1). Following collection, wing biopsy punches were placed into individually labelled tubes containing silica beads and stored at -20°C until DNA extraction. Although preservation protocols differed between study systems, experimental validation has shown no significant effect of storage method (liquid nitrogen vs silica bead preservation) on relative telomere length (rTL) estimates in bat wing tissue (57).

### (b) DNA extraction and relative telomere length measurement

Genomic DNA for both species was extracted from wing biopsies using membrane filter kits; Promega Wizard SV DNA Extraction kit (catalog no. A2371) for *R. ferrumequinum* or the Qiagen DNeasy Blood and Tissue Kit (Qiagen, CA USA) for *M. myotis*. Extractions carried out with the Promega kit were partially automated using a Hamilton STAR Deck liquid handling robot but otherwise followed manufacturer’s instructions. DNA concentration and purity were quantified for each sample using a BioDrop μLite Spectrophotometer (BioChrom, UK), with samples meeting a minimum 260/280 absorbance ratio of 1.7 retained for analysis. DNA integrity was confirmed for samples via 1% agarose gel electrophoresis. Relative telomere length (rTL) was measured by real-time quantitative PCR as the concentration of telomeric DNA relative to the mammalian brain-derived neurotrophic factor (BDNF) that is constant in number (single copy gene: SCG) and optimised for use in bats (57,70). Telomere and SCG reactions were carried out on separate plates due to differences in optimum annealing temperatures with reaction composition and cycling conditions following previously published protocols (57,69). Due to differences in the timing of data generation between the two study systems, qPCR procedures differed slightly between species. For *M. myotis*, telomere samples were assayed in triplicate on 384-well plates using the Applied Biosciences Quantstudio 7 Flex Real-Time PCR. Each plate was run with a negative control to detect contamination and a “golden sample” calibrator prepared from *M. myotis* wing tissue run in triplicate on each plate. For *R. ferrumequinum*, samples were assayed in triplicate on 96-well plates, and each plate layout was run twice for both telomere and SCG reactions on an Mx3000P qPCR system (Stratagene, CA, USA). A “golden sample” calibrator prepared from *R. ferrumequinum* wing tissue was included in triplicate on every plate to control for inter-plate variation, together with a negative control.

Raw data were exported from respective qPCR instruments and analysed using LinRegPCR (71) to perform baseline correction, calculating amplification efficiencies and quantification cycle (Cq) values (the cycle number at which the amplification curve crosses a fixed fluorescence threshold). For each species, telomere and single-copy gene data were normalised using the species-specific “golden sample” calibrator, extracted with the same kit as the experimental samples, to account for inter-plate variation and any systematic effects of extraction method (72). The average Cq was calculated across the six technical replicates for each sample for both amplicons for *R. ferrumequinum* and calculated across the three technical replicates for both amplicons for *M. myotis* respectively. rTL was calculated for each sample following (73) with further information on qPCR quality control detailed in Supplementary methods.

For *Myotis myotis*, average reaction efficiencies (mean ± SE) across the six qPCR plates were 2.004 ± 0.011 for the single-copy gene (SCG) reaction and 1.889 ± 0.004 for the telomere reaction. Intra-plate repeatability (intraclass correlation coefficient, ICC) calculated with the rptR package (Stoffel et al., 2017) was 0.909 ± 0.007 (95 % CI [0.895, 0.921], n = 1,488) for SCG Cq values and 0.848 ± 0.010 (95 % CI [0.825, 0.866], n = 1,488 technical replicates) for telomere Cq values. For *Rhinolophus ferrumequinum*, average reaction efficiencies (mean ± SE) across all plates were 1.968 ± 0.004 and 1.905 ± 0.007 for the single copy gene (SCG) and telomere reactions, respectively. The interplate repeatability (intraclass correlation coefficient), calculated using the reference sample (R package rptR, Stoffel et al., 2017), was 0.898 ± 0.015 (95% CI [0.866, 0.922], n = 384 samples, 124 plates) for SCG reactions and 0.808 ± 0.025 (95% CI [0.758, 0.852], n = 384 samples, 124 plates) for telomere reactions. Intraplate repeatability, calculated using all samples while controlling for plate effects, was 0.916 ± 0.004 (95% CI [0.909, 0.925]) for SCG Cq measures and 0.904 ± 0.005 (95% CI [0.897, 0.915], n = 5,238 technical replicates) for telomere Cq measures. While the above metrics confirm high assay repeatability, qPCR-based telomere assays do not distinguish terminal from interstitial telomeric sequences (ITS) and yield relative rather than absolute telomere length estimates (74,75). We assume that ITS content does not vary systematically with the predictors of interest within each species, while acknowledging that this assumption has not been directly tested, and that qPCR-based telomere measurements in bats have not been validated against an absolute method such as terminal restriction fragment (TRF) analysis.

To assess potential biases arising from tissue and DNA storage time, we fitted linear mixed-effects models separately for each species, with rTL as the response variable and tissue storage time (days) and DNA storage time (days) included as fixed effects. For *R. ferrumequinum*, age (in days) was additionally modelled as a quadratic term (see below). For both species, qPCR plate and year of sampling were included as random effects to account for technical and temporal variation, respectively. Individual identity was included as an additional random effect for *R. ferrumequinum*, as most individuals were sampled more than once. Neither tissue storage time (*M. myotis*: 95-3371 days; *R. ferrumequinum*: 17-666 days) nor DNA storage time (*M. myotis*: 106-112 days; *R. ferrumequinum*: 1-211 days) had a significant effect in either species (Supplementary Tables 2 and 3). Quadratic terms of both storage terms did not significantly improve models (*M. myotis*: LRT: χ2 = 1.846, *p* = 0.397; *R. ferrumequinum*: LRT: χ2 = 5.456, *p* = 0.062). Therefore, both variables were excluded from subsequent analyses.

### (c) Weather data collection

To quantify the influence of environmental conditions during development, climatic data were compiled for the regions surrounding the *M. myotis* and *R. ferrumequinum* maternity roosts. For Brittany, France, records of daily mean temperature (°C) and total daily rainfall (mm) were retrieved from Météo France (https://donneespubliques.meteofrance.fr/). The closest weather station to the roost site with consistent weather data for the ten years of sampling was the Arzal weather station (47◦52’ N, 2◦38’ W), located approximately 7 kilometres from the roost site. Given that *M. myotis* has a foraging range of several kilometres (76), this proximity makes the Arzal station a suitable proxy for the climatic conditions experienced by the study population. For Gloucestershire, UK, records of daily mean temperature (°C) and total daily rainfall (mm) were extracted from the Met Office Integrated Data Archive System (MIDAS; 77) from two weather stations active during this period approximately 8.1 (51◦41’ N, 2◦09’ W) and 12.2 km (51◦36’ N, 2◦13’ W) from Woodchester mansion, both within the range of *R. ferrumequinum* foraging area (54).

### (d) Statistical analyses

#### (i) Early development/Climwin

All statistical analyses were conducted in R 4.4.1 (78). rTL measurements were transformed to meet assumptions of normality and homoscedasticity prior to modelling. As rTL distributions differed between species, transformations were assessed separately for each species using Box-Cox procedures (79). A square-root transformation was applied to *M. myotis* rTL values, while a log transformation was applied to *R. ferrumequinum* rTL values. Following transformation, rTL values were standardised (z-scored) within species to facilitate comparison of effect sizes across models (80). All models were fitted using the R package *lme4* (81) unless otherwise specified. To assess whether early-life rTL varied significantly between annual birth cohorts, a linear mixed-effects model was fit with year included as a categorical factor for both species respectively. This approach allowed for direct comparison of mean rTL among year cohorts without assuming a continuous temporal trend. Both models included sex and scaled mass index (SMI) as fixed effects, and telomere assay plate as a random intercept to account for technical batch effects. SMI was calculated following (82), adjusting body mass to a common structural size using the scaling exponent derived from a log–log regression of mass on forearm length (see Supplementary Methods for full details). For *Rhinolophus ferrumequinum*, as exact age is known, and most were sampled more than once, age (in days) was included as a fixed effect and individual identity as a random intercept. As visual inspection suggested a non-linear relationship between rTL and age, an additional model including a quadratic age term was fitted. To determine if sexes differed in rTL as they aged, an interaction between sex and age was also tested.

As year effects were detected in both species (see Results), sliding window analyses were conducted with the *climwin* package (83). This approach systematically evaluates the effects of all possible climate windows on a response variable, enabling a data-driven assessment of when environmental conditions most strongly predict variation in TL. Because sampling regimes differed between species, climatic windows were defined in a manner appropriate to each system while maintaining a consistent analytical framework. In *M. myotis*, juveniles were sampled once annually at approximately 8 weeks of age, and climatic windows were therefore defined using an absolute time approach. A 60-day window preceding sampling was selected to encompass the postnatal growth and lactation period, during which telomere dynamics are most likely to be influenced by environmental conditions. In *R. ferrumequinum*, juveniles were sampled repeatedly within years, and climatic windows were defined relative to the timing of sampling. A 30-day window preceding each sampling event was used to capture short-term environmental exposure, approximating the typical interval between successive sampling occasions. Because many individuals were first sampled at, or within a few days of birth, the 30-day window preceding these initial sampling events primarily reflects late gestation and the immediate post-birth period experienced by mothers, rather than postnatal exposure of pups. For later sampling events, the same relative window increasingly captures postnatal environmental conditions experienced during early growth and the transition towards foraging independence. To improve interpretability, initial sampling events at birth within each year were treated as a single initial time point.

Within the specified windows for each species, *climwin* iteratively evaluated all possible sub-periods, testing the effects of temperature and precipitation using both linear and quadratic functions to allow for potential non-linear responses. In both species baseline models, sex and SMI were included as fixed effects, with year and qPCR plate fitted as random intercepts. Age in days modelled as a quadratic term was also fitted as a fixed term and individual identity as a random intercept in the *R. ferrumequinum* baseline model. Model support was assessed using ΔAICc, with more negative values indicating stronger support relative to the baseline model. To reduce the likelihood of biologically implausible or spurious associations, only climate windows spanning at least five consecutive days were considered biologically meaningful (61,83).

Statistical robustness was assessed using a randomisation procedure implemented in the randwin function, in which rTL data were permuted 100 times while maintaining the original climate structure. Two key metrics were used to assess the robustness of the climate window: (1) P_ΔAICc_, the proportion of randomised models with a ΔAICc lower than the observed best window, and (2) PC, the proportion of randomised models where the best climate window fell within the top 5% of ΔAICc values. A window was considered statistically supported if P_ΔAICc_ < 0.05 and PC < 0.50, ensuring that the identified climate effect was unlikely to have arisen by chance (83).

#### (ii) Maternal age

For *M.myotis*, maternal identity was determined through pedigree-based parentage assignments provided by collaborators as part of ongoing pedigree reconstruction efforts in this *M. myotis* population. These efforts build on the pedigree described in (59), which was developed using microsatellite genotyping. Maternal identity was assigned to 336 juveniles. Maternal age at the time of offspring birth was calculated based on the first time the female was caught and PIT tagged, utilizing long term mark-recapture data. Females first caught as juveniles had a known age, whereas those first captured as adults were assigned a minimum age of “1+” years. Maternal age at conception therefore represents the minimum known age of the mother at the time of conception throughout the *M. myotis* analyses.

Maternal identity in *R. ferrumequinum* was assigned based on field observations of pups attached to females during sampling, supplemented by the long-term pedigree for this population (84,85). Previous work in this population has established that attached pups are the female’s own offspring (65). Since 1982, intensive monitoring during the birth season (June-August) has ensured that nearly all pups are ringed within a few days of birth (54,64), providing exact birth dates and therefore known ages for many individuals. Maternal age at offspring birth was calculated directly from recorded birth dates where available.

To investigate the effect of maternal age at conception (MAC) on juvenile rTL, we fitted separate linear mixed-effects models (LMMs) for *M. myotis* and *R. ferrumequinum*. In each species, juvenile rTL z-scores were modelled as the response variable. Fixed effects included maternal age, offspring sex, and SMI. To assess sex-specific parental age effects, we included an interaction between maternal age and offspring sex. Random intercepts were included for telomere assay plate, sampling year, and maternal identity to account for batch effects, cohort effects, and non-independence among offspring from the same mother. To test if there were longitudinal patterns for maternal age, we partitioned maternal age into within- and between-mother components using within-subject centring (86). This approach separates within-mother ageing effects from between-mother differences, which may be influenced by selective disappearance (87). For each mother, the between-mother effect (mean maternal age) was calculated as the mean observed maternal age across all her sampled offspring. The within-mother effect (delta maternal age) was calculated as the deviation of maternal age for a given offspring from that mother’s mean age. We refitted the species-specific LMMs replacing the single maternal-age term with mean maternal age and delta maternal age, including interactions between each age component and offspring sex. To assess whether the sample size was sufficient to detect a biologically meaningful effect of maternal age for *M. myotis* and *R. ferrumequinum*, a simulation-based power analysis (500 simulations) was performed using the *simr* package (88).

#### (iii) Survival

To assess the association between rTL and next year survival, we used a bivariate modelling approach in a Bayesian framework using the R package *MCMCglmm* (89). For both species, rTL and first year survival (if an individual survived to the next summer) were fitted as joint response variables, with Gaussian and threshold distributions respectively. This approach enables joint estimation of variation in rTL and survival, as well as their covariance, at different hierarchical levels (90). For *M. myotis*, survival data was analysed for females only as most males do not return to the maternity colony after their first summer (91). Bats were assumed dead if they had not been recorded at the Férel roost or any other monitored sites for three consecutive years. Female *M. myotis* typically exhibit strong philopatry to maternity colonies, and this site is intensively monitored using fixed RFID antennas that provide continuous detection of PIT-tagged individuals. Although dispersal can occur, prolonged absence of greater than three years is considered to represent mortality in most cases. Survival estimates should therefore be interpreted as apparent survival within the monitored population. SMI was fitted as fixed effects for both response variables. Year sampled was fitted as a random effect for both variables, with qPCR plate fitted as a random effect for rTL only. Covariance at the year sampled level reflects associations between annual variation in survival and population level rTL, likely arising from shared environmental differences. In this model, residual covariance between traits captures phenotypic association such that positive covariance indicates a higher probability of survival among individuals with longer telomeres.

For *R. ferrumequinum*, first year survival was also analysed for females only, as survival estimates for immature males based on catch data are less reliable due to reduced and inconsistent recapture probability in subsequent years (54). Given the long-term and intensive monitoring of this colony, individuals not detected in their second year were assumed to have died. SMI was fitted as a fixed effect for both response variables. For rTL, age at sampling (in days) was included as a linear fixed effect, centred around the population mean, to facilitate interpretation of individual-level random intercepts. As rTL was measured repeatedly between individuals, whereas first year survival was measured once, survival was modelled as an individual-level trait. This allowed its variation to be part of the residual structure using the *covu* function in the prior (92,93). Individual identity was specified with an unstructured (co)variance matrix, enabling simultaneous estimation of among-individual variance in rTL and survival, as well as their covariance. By linking the random and residual effects structure, this allows survival to covary with both rTL and the slope of rTL change over time (93). Year was initially modelled with an unstructured covariance matrix however this term showed poor mixing of MCMC chains. Therefore, year and qPCR plate were fitted for rTL only.

## Results

### Year effects on early-life telomere length

#### Myotis myotis

Four out of the initial 500 samples failed DNA extractions and were excluded from the dataset, leaving a total of 496 individual juvenile *M. myotis* telomere samples (249 female and 247 male bats). We found a significant effect of birth year on early-life rTL (F_9,489_ = 22.726, *p* < 0.001) indicating strong interannual variation across cohorts (Figure 1A; Table S4). Sex (*β* = 0.084, *p* = 0.248, Figure 1C) and body condition (SMI; *β* = 0.007, *p* = 0.761; Table S4) were not significant predictors in this model.

**Figure 1.**
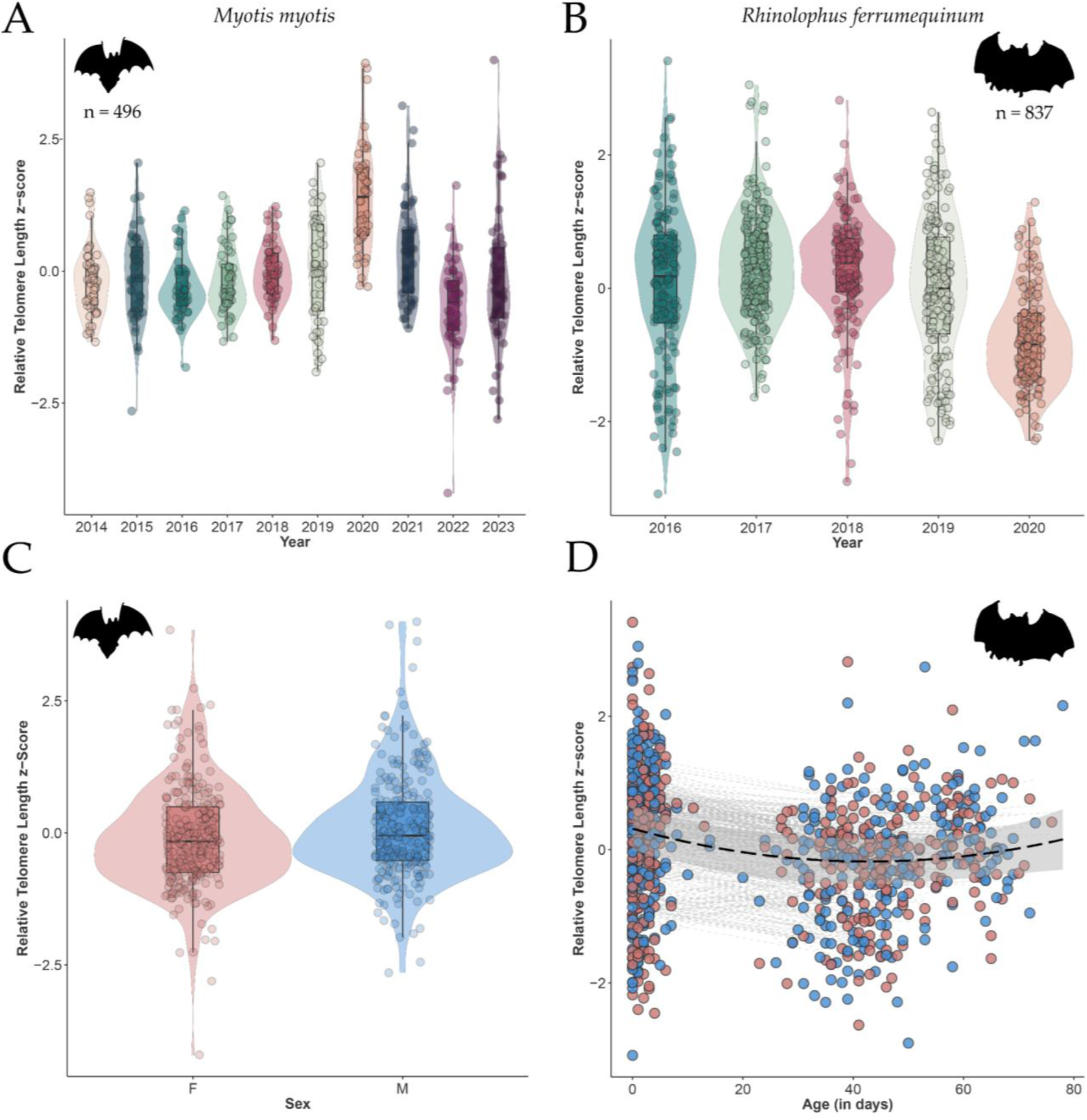
Early-life relative telomere length (rTL, z-scored) in juvenile *Myotis myotis* and *Rhinolophus ferrumequinum*. Panels A - B (top) show interannual variation in rTL across birth years for *M. myotis* (A) and *R. ferrumequinum* (B) with violin plots showing probability densities and overlaid data points (*M. myotis*, n = 496; *R. ferrumequinum*, n = 837). Box plots indicate medians and interquartile ranges. Panel C shows sex-specific distributions of rTL for *M. myotis* with violin plots of rTL z-scores for females (F, pink; *M. myotis*, n = 249) and males (M, blue; *M. myotis*, n = 247), where violin width reflects data density. Panel D shows *R. ferrumequinum* juvenile telomere dynamics during the early-life environment (0 - 78 days). Juveniles show an initial decline in telomere length with an increase occurring around 40 days, coinciding with the end of rapid growth. rTL data (z-scores) are colour coded by sex (Female: pink; Male: blue) with fitted lines based on model estimates with 95% confidence intervals.

#### Rhinolophus ferrumequinum

The *R. ferrumequinum* dataset comprised 837 relative telomere length (rTL) measurements from 435 individuals (213 females, 221 males). As in *M. myotis*, birth year had a significant effect on early-life rTL (F_4,396_ = *p* < 0.001; Figure 1B), whereas neither sex (*β* = 0.028, *p* = 0.587; Figure 1D) nor SMI (*β* = 0.042, *p* = 0.218, Table S5) showed a significant association. Age modelled as a quadratic term provided a better fit than a linear effect (ΔAICc = -6.47), revealing an initial decline in early-life rTL during the period of rapid juvenile growth (Figure 1D; Table S5). Notably, rTL increased from approximately 40 days of age onwards (Figure 1D), coinciding with the onset of flight and independent foraging. The interaction between age and sex was not significant (F₂,₅₈₁ = 0.867, *p* = 0.421), indicating that age-related rTL trajectories did not differ between males and females (Figure 1D).

### Sliding window analysis of climate effects

#### Myotis myotis

To refine the relationship between climate and rTL, a sliding window analysis was performed using a 60-day window prior to *M. myotis* sampling. This approach identified the period during which rainfall and temperature were most strongly associated with juvenile rTL. The best-supported model (ΔAICc = -19.29) indicated that mean daily precipitation within a window spanning 24 to 15 days before sampling was the strongest predictor of rTL, with a positive quadratic relationship providing the best fit (Figure 2, Table S6). This period of time corresponds to early- to mid-June, when *M. myotis* pups are approximately 5 - 6 weeks old. The final model assessing rainfall effects confirmed a significant quadratic relationship (*β* = 8.605, *p* < 0.001), with rTL increasing with rainfall but showing some non-linearity in response (Table S7). Because precipitation and temperature were strongly collinear within the identified windows, each variable was subsequently analysed in separate models. Temperature also showed a positive quadratic relationship (ΔAICc = -7.43; Table S6), with a window spanning 36-41 days prior to sampling, and longer rTL with higher temperatures (*β* = 6.679, *p* = 0.008; Table S8).

**Figure 2:**
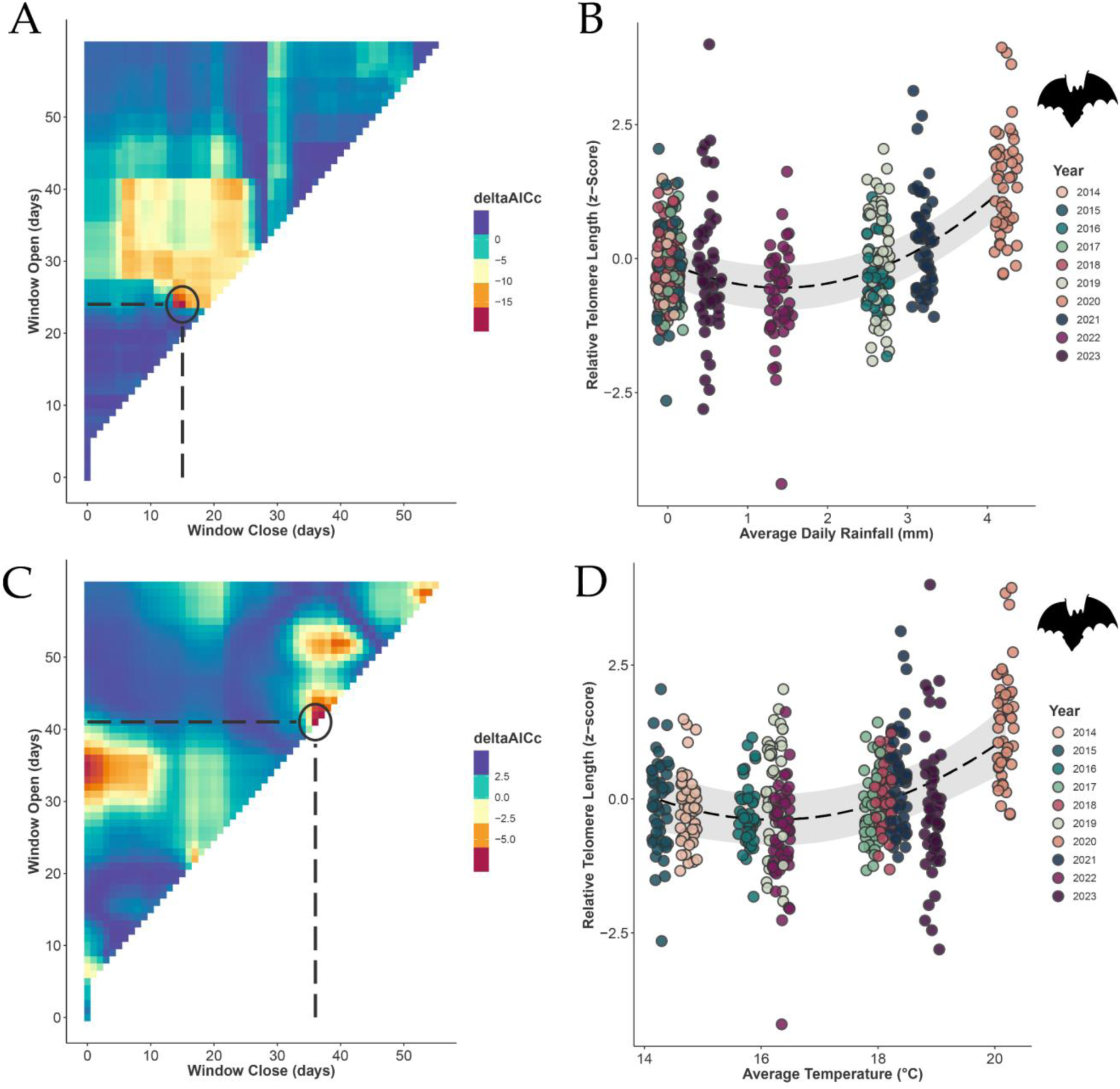
Sliding window analysis for the effect of average daily rainfall (mm) on early-life telomere length in *Myotis myotis* bats. Panels A and C: DeltaAICc values represent the differences between the AICc of each climate window model and the baseline model with no climate variable. Colours represent the deltaAICc values, with warmer colours indicating stronger effects of rainfall and average temperature on telomere length. Circle and dotted lines represent the most sensitive time window ≥ 5 days with results indicating the most significant window to be 15 to 24 days for rainfall and before sampling 36 to 41 days for temperature. Panel B: Relationship between average daily rainfall (during window specified from *climwin*) and relative telomere length (z-score) in juvenile bats across multiple years. Panel D: Relationship between average temperature (during window specified from *climwin*) and relative telomere length (z-score) in juvenile bats across multiple years. Dashed black lines represents the fitted quadratic relationship between telomere length and daily rainfall, with a 95% confidence interval shaded.

#### Rhinolophus ferrumequinum

The best-supported model for *R. ferrumequinum* (ΔAICc = -30.60) indicated that mean daily precipitation within a window spanning 30 to 25 days before sampling was the strongest predictor of rTL, with a quadratic relationship providing the best fit (Figure 3A, Table S6). Temperature also showed a quadratic relation (ΔAICc = -24.23), with a window spanning 30 to 15 days prior to sampling (Figure 3C, Table S6). As with *M. myotis*, weather variables were collinear and analysed in separate models. Both variables showed significant non-linear associations with juvenile rTL. Precipitation showed a significant positive quadratic relationship (*p* < 0.001), with rTL increasing with higher rainfall levels (Figure 3B, Table S9). In contrast to *M. myotis*, temperature exhibited a significant negative quadratic relationship (*β* = -6.094, *p* < 0.001), indicating an optimal range of environmental conditions results in longer early-life rTL (Figure 3D, Table S10). Randomisation tests for both species indicate that these signals are unlikely to reflect false positives, with PΔAICc < 0.05 and PC < 0.5 (Figure S1).

**Figure 3:**
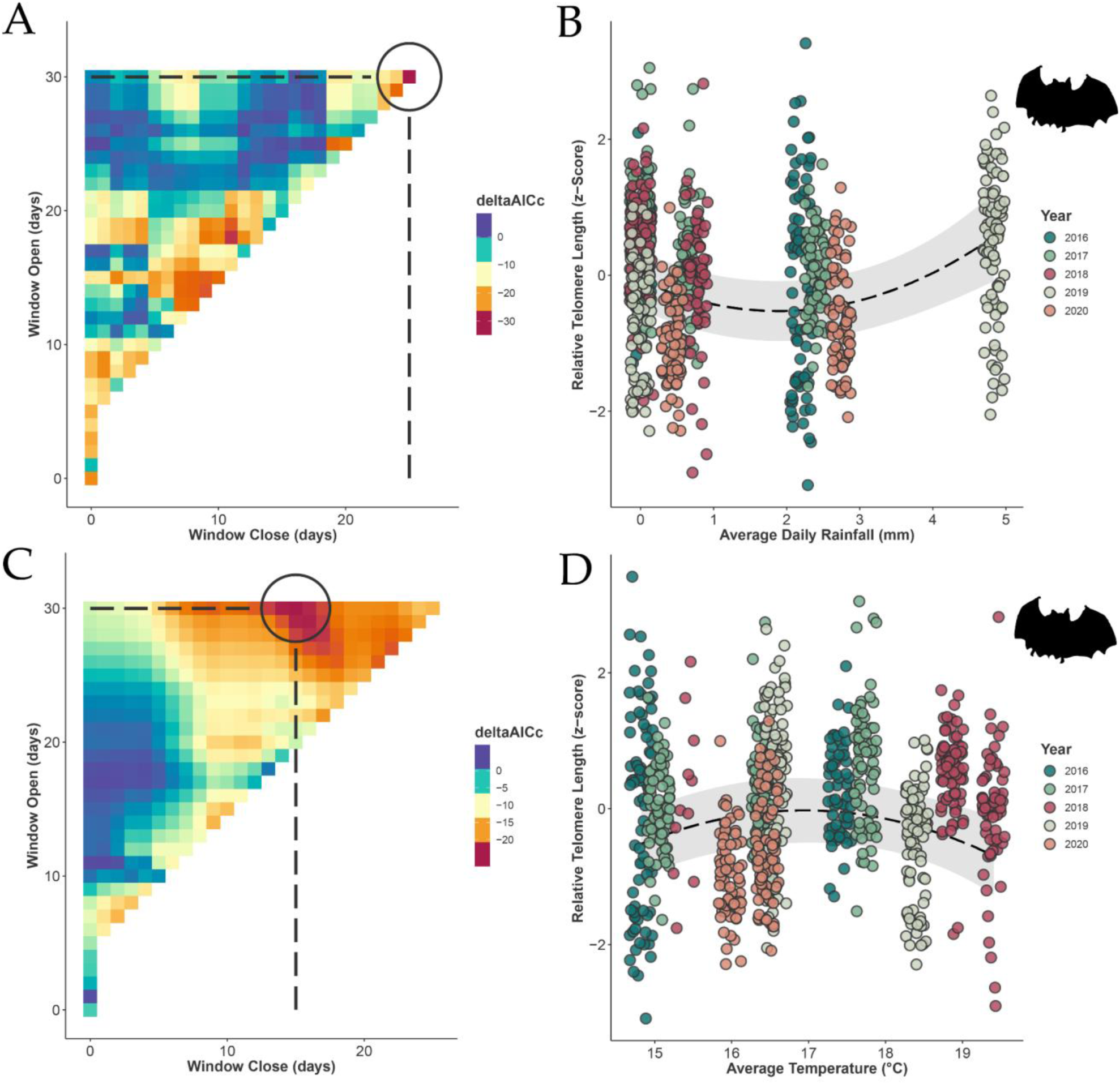
Sliding window analysis for the effect of average daily rainfall (mm) and average temperature on early-life telomere length in juvenile *Rhinolophus ferrumequinum*. Panels A and C: DeltaAICc values represent the differences between the AICc of each climate window model and the baseline model with no climate variable. Colours represent the deltaAICc values, with warmer colours indicating stronger effects of rainfall or average temperature on telomere length. Circle and dotted lines represent the most sensitive time window ≥ 5 days with results indicating the most significant window to be 25 to 30 days before sampling for rainfall and 15 to 30 days for temperature. Panel B: Relationship between average daily rainfall (during window specified from climwin) and relative telomere length (z-score) in *R. ferrumequinum*. Panel D: Relationship between average temperature (during window specified from climwin) and relative telomere length (z-score) in *R. ferrumequinum*. Dashed black lines represent model estimates with 95% confidence intervals from respective models.

### Maternal age effects

#### Myotis myotis

Of the dataset, 336 samples had known mothers, ranging from 1-11 years and were included in analyses of maternal age effects (Figure S2A). In an initial model including maternal age at conception (MAC) as a single fixed effect, there was no significant association between MAC and juvenile rTL (*β* = 0.051, *p* = 0.060, Figure 4A, Table S11), and no interaction between MAC and offspring sex (*β* = -0.043, *p* = 0.269, Table S11). To distinguish between population-level differences among mothers and longitudinal changes within the same mothers across years, MAC was then partitioned into a between-mother (mean MAC) and within-mother (delta MAC) component. Neither the between-mother effect (mean MAC; *β* = 0.039, *p* = 0.077) nor the within-mother effect (delta MAC; *β* = 0.056, *p* = 0.463) was significantly associated with juvenile rTL (Figure 4B, Table S12). There was no significant interaction between maternal age components and sex, indicating that maternal age effects did not differ between male and female juveniles (Table S12). Neither sex nor body condition (SMI) showed a significant association with juvenile rTL in this model. Simulation-based power analysis indicated that, accounting for model structure and sample size, there was ≥80% power to detect a maternal age at conception (MAC) effect of 0.055 or greater (Figure S2B). To facilitate comparison with previous studies, this effect size was converted to a correlation coefficient (e.g. MAC effect size*(SDMAC/SDrTL)) (40,94). This resulted in an estimated correlation coefficient of r = 0.086, suggesting that while the analysis had sufficient power to detect moderate maternal age effects, smaller effect sizes cannot be excluded.

**Figure 4.**
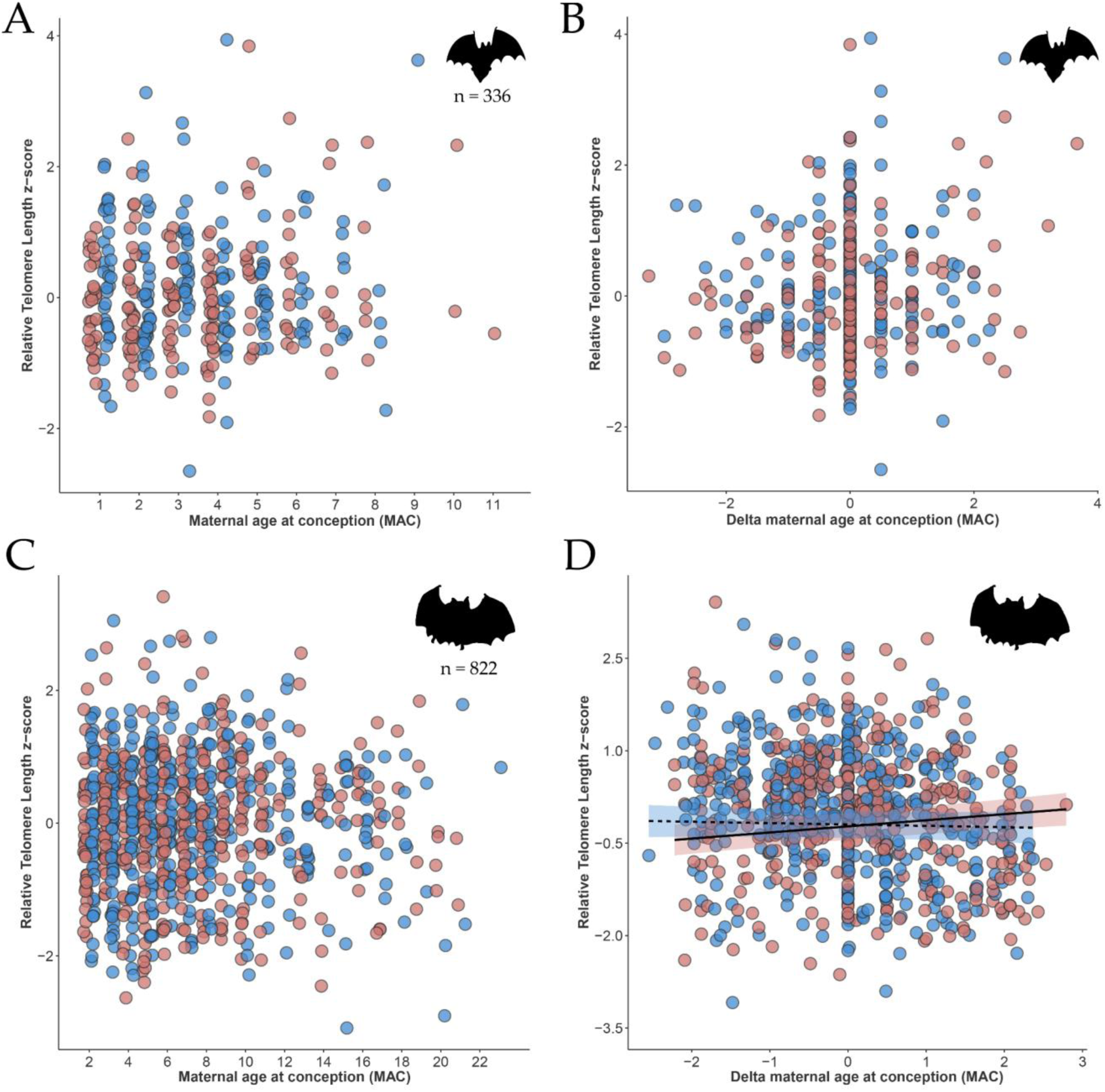
Maternal age effects on juvenile relative telomere length (rTL, z-score) in *Myotis myotis* and *Rhinolophus ferrumequinum*. Panels A and B: The relationship between maternal age at conception (MAC), within-mother maternal age at conception (delta MAC) and juvenile rTL (n = 336) in *M. myotis*. Neither MAC or partitioned age categories (mean MAC and delta MAC) were significant with no significant interactions with offspring sex (Table S11 and S12). Panels C and D: The relationship between MAC, delta MAC and juvenile rTL (n = 822) in *R. ferrumequinum*. While overall MAC was not significant, there was a significant interaction between offspring sex and delta MAC (F₁,₄₀₀ = 5.176, *p* = 0.024), indicating sex-specific longitudinal slopes. Points represent individual juveniles; fitted lines show model predictions with 95% confidence intervals (males: dotted; females: solid). Offspring sex is indicated by colour (males: blue; females: pink).

#### Rhinolophus ferrumequinum

For 421 juveniles (822 rTL samples), maternal age was known (age range 2-23 years old), representing offspring from 150 identified mothers (Figure S2C). In the single fixed effects model, there was no significant association between MAC and juvenile rTL (β = 0.002, p = 0.834), and no interaction between MAC and offspring sex (β = -0.004, p = 0.766; Figure 4C, Table S13). However, when considering longitudinal effects of MAC, daughters exhibited a positive within-mother age slope (*β* = 0.009) and sons a negative slope (*β* = -0.002), with neither slope significantly different from zero but significantly different from one another (*β* = -0.121, *p* = 0.024; Figure 4D, Table S14). Accounting for model structure and size, there was ≥80% power to detect MAC effect sizes ≥ 0.030, equivalent to a correlation coefficient of r = 0.131.

### First year survival

Using bivariate Bayesian mixed models, we examined covariation between rTL and survival to the subsequent summer in both species. In juvenile female *M. myotis*, we detected no significant covariance between rTL and survival at either the among-individual (σ = -0.013, 95% CI [-0.153, 0.124]) or among-year levels (σ = 0.139, 95% CI [-0.234, 0.941]; Supplementary Table S15). In *R. ferrumequinum* females, survival was also not associated with rTL (σ = -0.006, 95% CI [-0.055, 0.046]) and we found no evidence that individual rates of telomere change were related to survival (σ = -0.001, 95% CI [-0.010, 0.008]; Supplementary Table S16).

## Discussion

Here we examined early-life telomere dynamics in two long-lived, temperate bat species, *Myotis myotis* and *Rhinolophus ferrumequinum*, both of which exhibit exceptional longevity relative to body size using a unique dataset to enable us to explore early-life effects. We found that early-life rTL was shaped by short-term climatic conditions during the developmental window in both species, with non-linear effects of rainfall and temperature in *M. myotis* and *R. ferrumequinum*. Within-juvenile rTL in *R. ferrumequinum* followed a non-linear trajectory of decline followed by recovery as pups transitioned to independent foraging. Maternal age effects on offspring rTL were limited overall, with a weak sex-specific longitudinal effect detected only in *R. ferrumequinum* daughters. Juvenile rTL did not predict early survival in either species. Together, these findings provide the first detailed characterisation of early-life telomere variation in bats and identify climatic conditions during the developmental window as the most consistent driver of variation in juvenile rTL.

### Early life and climatic conditions on juvenile telomere length

In both species, the relationship between rainfall and juvenile rTL was best described by a positive quadratic function, with longer telomeres at both low and high rainfall levels. One plausible explanation is that resource limitation at either rainfall extreme slows juvenile growth, which in turn may reduce cellular proliferation and the accompanying telomere attrition during early development. Rapid early growth has been repeatedly associated with accelerated telomere loss across taxa (18), and conversely, conditions that constrain growth rate, through reduced maternal provisioning, delayed weaning, or slower accrual of body size, may result in longer telomeres at a given developmental stage. In *M. myotis*, which primarily consumes ground-dwelling arthropods such as carabid beetles (95), variation in rainfall may influence prey availability through effects on soil moisture and beetle activity (96,97), with consequences for lactational investment and juvenile growth trajectories. In *R. ferrumequinum*, rainfall is particularly important for the availability of dung- and soil-associated insects such as *Acrossus rufipes* which dominate juvenile diets during early independence, with both drought and waterlogging potentially suppressing larval development and adult beetle emergence (51,98). Selective disappearance of pups with shorter telomeres under adverse conditions at either rainfall extreme may additionally contribute to the observed pattern (23). The specific mechanism cannot be fully resolved with the current data, and the non-linear rainfall–rTL relationship observed here warrants further investigation, ideally with direct measurement of maternal foraging success and pup growth rate across the rainfall gradient.

Previous studies in these populations have examined the relationship between rainfall and telomere length at later life stages. Rainfall during the spring transition has been associated with longitudinal telomere shortening in adult *M. myotis* across a broader age range (59). Similarly (61) reported that cumulative rainfall during the hibernation period was associated with shorter telomeres in adult *R. ferrumequinum*, arguing for reduced aerial insect availability and increased arousal frequency from torpor under milder, wetter conditions. The apparent directional difference with our juvenile results is not a contradiction but likely a reflection of the different biological processes operating at different life stages, seasons, and temporal scales. In adults, rainfall during active foraging periods imposes direct energetic and thermoregulatory costs (59, 61), whereas in pups rainfall effects are mediated largely indirectly through maternal foraging success and milk provisioning, with direct exposure increasing only as juveniles begin independent foraging. The temporal scales also differ substantially: our windows span 15–30 days around the summer lactation and growth period, whereas (59) examined spring weather in relation to year-on-year changes in adult TL, and (61) examined cumulative rainfall across the full winter hibernation period. Taken together, these results suggest that the relationship between rainfall and telomere dynamics in bats is context-dependent, varying with life stage, season, and temporal scale, and that generalising rainfall effects across life stages risks obscuring biologically meaningful differences between adult physiological costs and juvenile developmental processes.

Temperature effects were observed in both species, though the direction of these relationships differed. In *M. myotis*, juvenile rTL showed a positive quadratic relationship with temperature, with longer rTL associated with warmer conditions during the critical window 36-41 days prior to sampling. This window corresponds to the early postnatal period in late May to early June. During this period, pups are dependent on maternal milk, and lactating females experience decreased thermoregulatory costs at higher ambient temperatures, allowing greater energy allocation to milk production (99). As pups develop, warmer conditions may additionally support greater insect availability and foraging efficiency for both mothers and increasingly independent juveniles (99). Together, these suggest that warmer early postnatal conditions improve the nutritional environment during a critical developmental period, enabling greater investment in somatic maintenance reflected in longer rTL (18). This is broadly consistent with findings in the same population showing positive associations between rTL and temperature across a wider age range (59).

By contrast, *R. ferrumequinum* exhibited a negative quadratic relationship between temperature and juvenile rTL with shorter telomeres at higher temperatures. This pattern suggests that intermediate thermal conditions are most favourable for early-life telomere maintenance, while elevated temperatures may impose indirect developmental costs. These likely arise through temperature and moisture driven constraints on prey phenology. Juvenile *R. ferrumequinum* do not forage independently until approximately 30 days of age and rely heavily on seasonally abundant prey such as the night flying dung beetle *Acrossus rufipes*. Drought conditions can delay peak activity of this prey until later in the season (98), potentially creating a phenological mismatch between juvenile energetic demand and food availability if warm spring conditions lead to earlier births (55). Similar sensitivity of bats to adverse or extreme temperature conditions have been documented in other long-lived bat species. For example, mass mortalities have been recorded in flying-foxes (*Pteropus poliocephalus*) during summers that reach extremely hot temperatures (100,101) and low juvenile survival has been observed in big brown bats (*Eptesicus fuscus*) when born during particularly warm and dry years (102).

### Within-individual telomere dynamics during early development

Within-juvenile telomere changes in *R. ferrumequinum* followed a non-linear trajectory, with rTL initially declining during the rapid growth phase and then increasing from approximately 40 days of age onwards. This pattern is consistent with the growth-associated telomere attrition documented across vertebrates (18) and reflects the period of rapid postnatal development in this species, during which pups transition from exclusive maternal provisioning (until ∼30 days of age) to increasingly independent foraging while still completing skeletal and echolocation development (50,51,103). The subsequent increase in rTL, coinciding with the onset of flight and independent foraging, suggests that active telomere maintenance or lengthening mechanisms may be engaged following the intense early-growth period. Such non-linear early-life trajectories, with initial decline followed by recovery, have not previously been documented in juvenile bats of other species, although cross-sectional data from *R. ferrumequinum* 0 - 1 year-olds suggested an apparent lengthening pattern (57). Our longitudinal data refines this pattern, showing that rTL does not simply lengthen across the first year, but instead declines during peak growth before increasing as pups become nutritionally independent. These findings also complement longitudinal work in *R. ferrumequinum*, where within-individual increases in rTL have been observed during hibernation (60) alongside evidence of increased telomerase expression. This suggests that *R. ferrumequinum* is capable of telomere maintenance or lengthening at multiple life stages, including during the post-growth period of early independence and during winter torpor, despite the overall pattern of age-related telomere shortening documented across this species’ lifespan (57). We note that comparable longitudinal early-life telomere data are not available for other bat species, and that within-individual telomere trajectories during the juvenile growth period represent a gap in the broader bat telomere literature.

### Maternal age effects on offspring telomere length

Despite the potential importance of paternal germline mechanisms highlighted in some taxa, our analyses focused on maternal age at conception, reflecting both the structure of the study systems as maternity colonies and the biological relevance of maternal investment during early development in bats. Maternal identity and age are reliably known through long-term monitoring, whereas paternity assignment remains challenging in these systems (largely due to difficulties in locating and sampling breeding males), precluding robust inference on paternal age effects. In *M. myotis*, maternal age at conception was not significantly associated with juvenile rTL at either the population (between-mother) or longitudinal (within-mother) level, and no sex-specific effects were detected. Power analyses indicated that the study was sufficiently powerful to detect moderate maternal age effects, but smaller effects cannot be excluded. Several factors may explain the absence of maternal age effects in this study. The relatively young and narrow maternal age range observed (1-11 years), given this species’ potential lifespan (37 years), may have limited the ability to detect strong age-related effects. Similarly, the telomere maintenance previously documented in adult *M. myotis* over the sampled age-range (0-7 years old; 57) may indicate limited telomere attrition during early adulthood, potentially reducing the expected maternal age effects that arise from telomere attrition observed in other taxa. Similar absences of parental age effects on offspring TL have been reported in other mammals, including Soay sheep and European badgers (40,104).

In *Rhinolophus ferrumequinum*, maternal age effects were absent when assessed cross-sectionally. However, partitioning maternal age into within- and between-mother components revealed a small but significant sex-specific longitudinal effect, where daughters compared to sons born to older mothers exhibited slightly longer telomeres. The mechanisms by which or why sex-specific parental age at conception effects occur are unclear. Suggestions include differences in how sons and daughters respond to parental environments or sex-specific inheritance or investment (43,44). In *R. ferrumequinum*, females are the larger sex and exhibit higher first year survival probabilities, with sex ratios in this population known to shift toward males following prolonged periods of poor weather (55). Together, these patterns suggest that older mothers may favour investment in daughters under favourable conditions, with potential implications for early-life telomere dynamics.

Overall, our findings indicate that maternal age effects on offspring TL in bats are generally weak, with subtle sex-specific patterns emerging only when longitudinal maternal age is considered. These results add to growing evidence that parental age effects on telomeres are best resolved using longitudinal studies. However, it is important to note that maternal age at conception may not fully capture cumulative reproductive effort. Females of the same chronological age can differ substantially in birth sequence and lifetime reproductive history, particularly in long-lived species where reproductive skipping and variable annual investment occur (60,105). Such variation could influence offspring condition. Future work integrating detailed reproductive histories for both species will be necessary to disentangle maternal age effects from cumulative reproductive stress in bats. Furthermore, maternal age effects may not be linear across the lifespan. If reproductive senescence occurs only at advanced ages in these long-lived bats, age-related influences on offspring TL may emerge only beyond the age range represented in our dataset. Consequently, the absence of strong or consistent associations in this study does not exclude the possibility of late-life maternal age effects.

### Early-life telomere length and survival

Despite evidence that early-life TL was shaped by environmental conditions, we found no support for early-life rTL predicting short-term survival in either species. These findings are consistent with studies suggesting that TL is not a universal predictor of survival (e.g. 106,107), despite positive associations reported in some taxa (e.g. 23,34). One likely explanation relates to the unusual ageing biology of bats. Compared to other mammals of similar body size, bats exhibit exceptional longevity and, in many species, delayed senescence, supported by mechanisms linked to cellular maintenance, oxidative stress resistance, and genome stability (47–49,56,108,109). In *M. myotis*, adult rTL shows little or no decline with age (57,59), suggesting that telomere dynamics do not follow the progressive attrition patterns observed in many vertebrates. This stability implies evolved mechanisms of telomere maintenance or repair, potentially involving sustained telomerase activity or alternative telomere lengthening pathways, therefore early-life telomere variation may be less consequential in *Myotis*. The lack of a significant correlation between early-life TL and survival to first year in both species may also be attributed to hibernation. Hibernation induces a state of torpor characterized by reduced metabolic activity and body temperature (110), which can mitigate oxidative stress and slow cellular ageing processes (111). Given evidence of slower telomere shortening rates during hibernation in the same population of *R. ferrumequinum*, (61), this physiological adaptation may buffer individuals against the detrimental impacts of shorter early-life telomeres, thereby decoupling the relationship between juvenile TL and subsequent survival. These buffering processes may be particularly effective in *M. myotis*, where TL remains stable across adulthood, compared to *R. ferrumequinum*, where age-related shortening is evident (61). Consequently, early-life variation in TL may have weaker fitness consequences in species with stronger telomere maintenance capacity.

Several limitations should be considered when interpreting these results. In both species, survival analyses focused on first-year survival, which captures only an early and relatively short-term fitness component and may not reflect longer-term consequences of early-life telomere variation. Additionally, analyses were restricted to females, reflecting the structure of the maternity roosts studied, and therefore cannot exclude sex-specific survival effects. Furthermore, in *M. myotis*, early-life TL was measured at a single time point, preventing direct estimation of telomere attrition during early development. By contrast, repeated early-life sampling in some *R. ferrumequinum* individuals allowed us to estimate short-term telomere change during development and test whether rates of attrition, rather than absolute TL, predicted first-year survival. Evidence from other wild systems suggests that rates of telomere change, rather than TL measured at a single time point, can be more informative predictors of survival (e.g. 112). Despite this, we found no evidence that early-life telomere change predicted first-year survival in *R. ferrumequinum.* Consequently, early-life telomere dynamics may influence later-life performance or survival in bats through delayed or cumulative effects that are not detectable within the scope of first-year survival analyses. Longer-term monitoring with repeated sampling across development and adulthood will be essential to evaluate these possibilities in long-lived, hibernating mammals.

## Conclusion

Overall, our findings show that variation in juvenile TL in bats is shaped by short-term weather conditions experienced during early development. Rainfall predicted juvenile rTL in both species through a non-linear relationship, while temperature effects diverged: warmer conditions were associated with longer rTL in *M. myotis*, whereas intermediate temperatures were most favourable in *R. ferrumequinum*. In contrast, maternal age at conception had little influence on offspring TL in either species, and early-life TL did not predict short-term survival. Together, these results suggest that in long-lived, hibernating bats, early-life telomere variation primarily reflects developmental environmental conditions rather than parental age or immediate survival prospects. These patterns are consistent with the exceptional physiological adaptation of bats, including effective telomere maintenance and physiological buffering through hibernation. Such adaptations may reduce any potential longer-term consequences of environmentally driven telomere variation early in life. By providing the first insight into early-life telomere dynamics in bats, our results emphasise that the biological impact of TL depends on species ecology and temporal scale, underscoring the need for longitudinal and life-history-informed approaches when using telomeres as biomarkers of ageing and fitness in natural systems.

## Acknowledgments

For assistance in the *Myotis myotis* fieldwork, we would like to thank all the members of Bretagne Vivante, students and volunteers from University College Dublin, and all owners/local authorities for allowing access to the sites. We gratefully acknowledge the Woodchester Mansion Trust for access to the *Rhinolophus ferrumequinum* maternity colony and the many volunteers past and present who have helped to collect *R. ferrumequinum* data over the years.

## Ethics

*Rhinolophus ferrumequinum*: All *Rhinolophus ferrumequinum* bats were caught and sampled under licenses (Natural England 2015-11974-SCI-SCI; 2016-25216-SCI-SCI; 2017-31148-SCI-SCI) issued to G.J., with tissue biopsy procedures additionally licensed under Home Office Project Licenses (PPL 30/3025 prior to 2018; P307F1428 from 2018 onward) and Home Office personal licenses.

*Myotis myotis*: Sampling was carried out in accordance with sampling and ethical guidelines as detailed in successive capture and processing permits issued to F.T. and S.J.P. by the Prefet de Morbihan (Brittany) for the duration of the study period (PROGCACCHI_007). Access to all field sites was granted by local authorities in collaboration with Bretagne Vivante.

## Supplementary material

Supplementary materials have been submitted with the main text.

## Data and code availability

The datasets and code supporting this article will be available with supplementary material upon acceptance.

