## Supplementary Materials for "Early-life telomere length variation under changing developmental conditions in long-lived bats"

**Supplementary methods**

To amplify telomere sequences, primers tel1 (5′-GGTTTTTGAGGGTGAGGGTGAGGGTGAGGGTGAGGGT-3′) and tel2 (5′-TCCCGACTATCCCTATCCCTATCCCTATCCCTATCC-3′) were used (1). The single copy gene, BDNF (mammalian brain-derived neurotrophic factor), was amplified using primers BDNF_F1 (5′-AGCTGAGCGTATGTGACAGT-3′) and BDNF_R1 (5′-TGGGATTACACTTGGTCTCGT-3′) from (2). To minimise potential confounding between qPCR plate effects and biological variables such as sampling year, *Myotis myotis* samples were organised using a slicing approach (3). This method ensures samples from different years were deliberately distributed across plates rather than grouped sequentially or randomly. By "slicing" across sampling years, this allows for the statistical separation of technical variation (e.g., plate effects) from biological variation such as cohort or environmental effects.

For *Rhinolophus ferrumequinum,* samples were randomly allocated onto extractions plates along with other samples collected as part of this long-term study (4,5). Although randomisation minimises confounding between biological variables (e.g. sampling year) and batch effects, residual between-plate variation can occur in qPCR assays (3). Therefore, qPCR plate identity was included as a covariate in all statistical models to account for technical variance.

For both species, reactions were carried out in 10μl volumes with 5μl of Takara SYBR Green (Clontech), 0.2μl of ROX, 2.5μl of DNA with the remaining volume made up with PCR grade water (Invitrogen). Telomere and SCG reactions were the same except for primer volumes with all primers used at 50μM concentration (0.027μl tel1, 0.09μl tel2, 0.12μl BDNF_F1, 0.2μl BDNF_R1). For baseline correction, calculating amplification efficiencies and Cq values (the number of cycles the qPCR amplification curve crosses a set fluorescence threshold), the software package LinRegPCR was used (6). After normalization using the golden sample as reference, the average Cq was calculated across the replicates for each sample for both amplicons. rTL was calculated for each sample, following (7):

$$rTL = \frac{(E_{TEL})^{Cq_{TEL(Calibrator)} - Cq_{TEL(Sample)}}}{(E_{SCG})^{Cq_{SCG(Calibrator)} - Cq_{SCG(Sample)}}}$$

Where $E_{TEL}$ and $E_{SCG}$ are the amplicon and plate specific qPCR efficiencies, $Cq_{TEL(Calibrator)}$ and $Cq_{SCG(Calibrator)}$ are the average Cq values for the “golden sample” calibrator for each amplicon and $Cq_{TEL(Sample)}$ and $Cq_{SCG(Sample)}$ are the average Cq values for individual samples for each amplicon respectively.

**Scaled Mass Index**

For both species, models included scaled mass index (SMI) as a fixed effect. SMI was calculated following (8) as:

$$SMI = W_i \left(\frac{L_0}{L_i}\right) b$$

where $W_i$ is body mass, $L_i$ is forearm length (as a measure of structural size), $L_0$ is the mean forearm length of the sampled population, and *b* is the scaling exponent estimated from an ordinary least squares regression of log-transformed mass on log-transformed forearm length. Forearm length was used as the linear body size metric.

**Table S1: Summary of the *Rhinolophus ferrumequinum* samples caught during early-life between 2016 and 2020 in the Woodchester Mansion population. There were 847 rTL measurements taken from 441 individuals.**

| *Rhinolophus ferrumequinum* | | | |
|---|---|---|---|
| | **Number of samples** | **Number of individuals** | |
| | **1** | 72 | |
| | **2** | 306 | |
| | **3** | 63 | |
| **Number of samples by year** | | **Number of samples by month** | |
| **2016** | 149 | **June** | 100 |
| **2017** | 233 | **July** | 333 |
| **2018** | 164 | **August** | 293 |
| **2019** | 172 | **September** | 147 |
| **2020** | 155 | | |

**Table S2: Summary of mixed-effects model testing the effect of sample storage time (days) and DNA storage time (days) on *Myotis myotis* rTL.** Significant values ($p < 0.05$) are in bold.

| Effects | Estimate | Std Error | t value | p-value |
|---|---|---|---|---|
| **Fixed Effects** | | | | |
| (Intercept) | 0.072 | 0.265 | 0.270 | 0.794 |
| Sample storage time | 0.010 | 0.165 | 0.058 | 0.954 |
| DNA storage time | -0.063 | 0.079 | -0.803 | 0.423 |
| **Random Effects** | **Variance** | | | |
| Year | 0.257 | | | |
| qPCR Plate | 0.234 | | | |
| Residual | 0.599 | | | |

**Table S3: Summary of mixed-effects model testing the effect sample storage time (days) and DNA storage time (days) on *Rhinolophus ferrumequinum* rTL.** Significant values ($p < 0.05$) are in bold.

| Effects | Estimate | Std Error | t value | p-value |
|---|---|---|---|---|
| **Fixed Effects** | | | | |
| (Intercept) | -0.061 | 0.165 | -0.367 | 0.781 |
| Age (in days) | -0.187 | 0.033 | -5.731 | <0.001 |
| Age (in days)$^2$ | 0.103 | 0.034 | 3.039 | 0.003 |
| Sample storage time | 0.089 | 0.087 | 1.022 | 0.311 |
| DNA storage time | -0.045 | 0.047 | -0.954 | 0.341 |
| **Random Effects** | **Variance** | | | |
| Year | 0.106 | | | |
| qPCR Plate | 0.261 | | | |
| Individual identity | 0.097 | | | |
| Residual | 0.479 | | | |

**Table S4: Summary of mixed-effects model testing the effect of birth year cohort (factor), sex, and scaled mass index (SMI) on juvenile telomere length (z-scores) in *Myotis myotis*.** Reference year is 2014. Significant values ($p$ < 0.05) are in bold.

| Effects | Estimate | Std Error | t value | p-value |
|---|---|---|---|---|
| **Fixed Effects** | | | | |
| (Intercept, Year 2014) | -0.239 | 0.515 | -0.464 | 0.644 |
| Year 2015 | -0.012 | 0.154 | -0.078 | 0.938 |
| Year 2016 | 0.018 | 0.156 | 0.118 | 0.907 |
| Year 2017 | 0.029 | 0.159 | 0.182 | 0.855 |
| Year 2018 | 0.196 | 0.155 | 1.260 | 0.208 |
| Year 2019 | 0.098 | 0.162 | 0.608 | 0.544 |
| **Year 2020** | **1.422** | **0.164** | **8.699** | **<0.001** |
| Year 2021 | 0.281 | 0.166 | 1.695 | 0.091 |
| **Year 2022** | **-0.761** | **0.179** | **-4.240** | **<0.001** |
| Year 2023 | -0.077 | 0.183 | -0.417 | 0.677 |
| Sex (Male) | 0.084 | 0.073 | 1.157 | 0.248 |
| SMI | 0.007 | 0.022 | 0.304 | 0.761 |
| **Random Effects** | **Variance** | | | |
| qPCR Plate | 0.238 | | | |
| Residual | 0.586 | | | |

**Table S5: Summary of mixed-effects model testing the effect of birth year cohort (factor), sex, age in days (quadratic function) and scaled mass index (SMI) on juvenile telomere length in *Rhinolophus ferrumequinum* (z-scores).** Reference year is 2016. Significant values ($p < 0.05$) are in bold.

| Effects | Estimate | Std Error | t value | p-value |
|---|---|---|---|---|
| **Fixed Effects** | | | | |
| (Intercept, Year 2016) | -0.463 | 0.457 | -1.013 | 0.311 |
| Year 2017 | 0.341 | 0.210 | 1.619 | 0.110 |
| Year 2018 | 0.032 | 0.223 | 0.144 | 0.886 |
| Year 2019 | -0.149 | 0.231 | -0.644 | 0.522 |
| **Year 2020** | **-0.946** | **0.250** | **-3.789** | **<0.001** |
| Sex (Male) | 0.028 | 0.059 | 0.474 | 0.636 |
| SMI | 0.042 | 0.034 | 1.233 | 0.218 |
| **Age (in days)** | **-5.375** | **0.940** | **-5.721** | **<0.001** |
| **Age (in days)$^2$** | **2.880** | **0.978** | **2.945** | **0.003** |
| **Random Effects** | **Variance** | | | |
| qPCR Plate | 0.236 | | | |
| Individual identity | 0.093 | | | |
| Residual | 0.481 | | | |

**Table S6: Summary of sliding window analysis for early-life rTL in relation to climatic variables (temperature and rainfall) for *Myotis myotis* and *Rhinolophus ferrumequinum*.** Different statistical functions (linear or quadratic) were used to model the relationships, and ΔAICc values indicate the relative model support compared to the null model. "WindowOpen" and "WindowClose" represent the days before sampling where climate variables were measured within a selected sliding window.

| | | | | | | |
|---|---|---|---|---|---|---|
| ***Myotis myotis*** | | | | | | |
| **Response variable** | **Climate** | **stat** | **func** | **DeltaAICc** | **WindowOpen** | **WindowClose** |
| Juvenile rTL | Temp | mean | lin | -5.92 | 59 | 54 |
| | Rain | mean | lin | -11.84 | 28 | 18 |
| | Temp | mean | quad | -7.43 | 41 | 36 |
| | Rain | mean | quad | -19.29 | 24 | 15 |
| ***Rhinolophus ferrumequinum*** | | | | | | |
| Juvenile rTL | Temp | mean | lin | -17.75 | 28 | 21 |
| | Rain | mean | lin | -25.10 | 8 | 1 |
| | Temp | mean | quad | -24.23 | 30 | 15 |
| | Rain | mean | quad | -30.60 | 30 | 25 |

**Table S7: Results of the linear mixed-effects models analysing the relationship between juvenile rTL in *M. myotis* and rainfall during the window identified by the *climwin* sliding window analysis.** Model includes sex and scaled mass index (SMI) as fixed effects, with qPCR plate treated as a random effect. Significant p-values (<0.05) are highlighted in bold.

| Effects | Estimate | Std Error | t value | p-value |
|---|---|---|---|---|
| **Fixed effects** | | | | |
| (Intercept) | -0.352 | 0.474 | -0.743 | 0.460 |
| Sex (Male) | 0.080 | 0.073 | 1.092 | 0.276 |
| Scaled Mass Index (SMI) | 0.016 | 0.020 | 0.788 | 0.432 |
| **Rainfall** | **6.471** | **1.041** | **6.219** | **<0.001** |
| **Rainfall$^2$** | **8.605** | **1.034** | **8.320** | **<0.001** |
| **Random Effects** | **Variance** | | | |
| qPCR Plate | 0.201 | | | |
| Year | 0.009 | | | |
| Residual | 0.598 | | | |

**Table S8: Results of the linear mixed-effects models analysing the relationship between juvenile rTL in *M. myotis* and temperature during the window identified by the *climwin* sliding window analysis.** The model includes sex and scaled mass index (SMI) as fixed effects, with qPCR plate treated as a random effect. Significant p-values (<0.05) are highlighted in bold.

| Effects | Estimate | Std Error | t value | p-value |
|---|---|---|---|---|
| **Fixed effects** | | | | |
| (Intercept) | -0.102 | 0.511 | -0.200 | 0.841 |
| Sex (Male) | 0.090 | 0.073 | 1.227 | 0.220 |
| Scaled Mass Index (SMI) | 0.005 | 0.022 | 0.214 | 0.831 |
| **Average Temperature** | **6.812** | **1.977** | **3.446** | **0.008** |
| **Average Temperature$^2$** | **6.679** | **1.938** | **3.446** | **0.008** |
| **Random Effects** | **Variance** | | | |
| qPCR Plate | 0.233 | | | |
| Year | 0.064 | | | |
| Residual | 0.598 | | | |

**Table S9: Results of the linear mixed-effects models analysing the relationship between juvenile rTL in *R. ferrumequinum* and rainfall during the window identified by the *climwin* sliding window analysis.** The model includes age in days (modelled as a quadratic term), sex and scaled mass index (SMI) as fixed effects, with qPCR plate, year sampled and individual identity treated as a random effect. Significant p-values (<0.05) are highlighted in bold.

| Effects | Estimate | Std Error | t value | p-value |
|---|---|---|---|---|
| **Fixed effects** | | | | |
| (Intercept) | -0.602 | 0.468 | -1.288 | 0.201 |
| **Age (in days)** | **-4.223** | **0.972** | **-4.344** | **<0.001** |
| Age (in days)$^2$ | 1.969 | 1.080 | 1.823 | 0.068 |
| Sex (Male) | 0.033 | 0.058 | 0.562 | 0.575 |
| Scaled Mass Index (SMI) | 0.042 | 0.033 | 1.269 | 0.205 |
| Rainfall | 2.976 | 1.864 | 1.596 | 0.112 |
| **Rainfall$^2$** | **8.313** | **1.590** | **5.230** | **<0.001** |
| **Random Effects** | **Variance** | | | |
| qPCR Plate | 0.247 | | | |
| Individual identity | 0.100 | | | |
| Year | 0.195 | | | |
| Residual | 0.453 | | | |

**Table S10: Results of the linear mixed-effects models analysing the relationship between juvenile rTL in *R. ferrumequinum* and temperature during the window identified by the climwin sliding window analysis.** Model includes age in days (modelled as a quadratic term), sex and scaled mass index (SMI) as fixed effects, with qPCR plate, year sampled and individual identity treated as a random effect. Significant p-values (<0.05) are highlighted in bold.

| Effects | Estimate | Std Error | t value | p-value |
|---|---|---|---|---|
| **Fixed effects** | | | | |
| (Intercept) | -0.745 | 0.477 | -1.562 | 0.122 |
| **Age (in days)** | **-5.640** | **0.926** | **-6.093** | **<0.001** |
| **Age (in days)$^2$** | **3.499** | **0.989** | **3.538** | **<0.001** |
| Sex (Male) | 0.029 | 0.058 | 0.497 | 0.619 |
| Scaled Mass Index (SMI) | 0.055 | 0.034 | 1.633 | 0.103 |
| Average Temperature | -1.102 | 1.828 | -0.603 | 0.547 |
| **Average Temperature$^2$** | **-6.094** | **1.480** | **-4.117** | **<0.001** |
| **Random Effects** | **Variance** | | | |
| qPCR Plate | 0.250 | | | |
| Individual identity | 0.097 | | | |
| Year | 0.229 | | | |
| Residual | 0.598 | | | |

**Table S11: Results of the linear mixed-effects model investigating the relationship between early-life rTL of *Myotis myotis* juveniles and maternal age at conception (MAC).** Significant values ($p<0.05$) are highlighted in bold.

| Effects | Estimate | Std Error | t value | p-value |
|---|---|---|---|---|
| **Fixed Effects** | | | | |
| (Intercept) | -0.262 | 0.573 | -0.457 | 0.648 |
| MAC | 0.051 | 0.027 | 1.872 | 0.062 |
| Sex (Male) | 0.173 | 0.156 | 1.106 | 0.270 |
| Scaled Mass Index | 0.011 | 0.025 | 0.454 | 0.650 |
| MAC * Sex (Male) | -0.042 | 0.038 | -1.107 | 0.269 |
| **Random Effects** | **Variance** | | | |
| Mother identity | 0.056 | | | |
| Year | 0.181 | | | |
| qPCR Plate | 0.214 | | | |
| Residual | 0.443 | | | |

**Table S12: Summary of mixed-effects model testing between and within mother at conception effects in *Myotis myotis*.** Significant values ($p<0.05$) are highlighted in bold.

| Effects | Estimate | Std Error | t value | p-value |
|---|---|---|---|---|
| **Fixed Effects** | | | | |
| (Intercept) | -0.299 | 0.581 | -0.514 | 0.608 |
| Mean MAC | 0.055 | 0.031 | 1.775 | 0.077 |
| Delta MAC | 0.038 | 0.054 | 0.699 | 0.485 |
| Sex (Male) | 0.156 | 0.174 | 0.898 | 0.370 |
| SMI | 0.012 | 0.025 | 0.501 | 0.617 |
| Sex (Male)*Mean MAC | -0.038 | 0.044 | -0.858 | 0.392 |
| Sex (Male)*Delta MAC | -0.054 | 0.076 | -0.714 | 0.476 |
| **Random Effects** | **Variance** | | | |
| qPCR Plate | 0.215 | | | |
| Mother identity | 0.054 | | | |
| Year | 0.187 | | | |
| Residual | 0.444 | | | |

**Table S13: Results of the linear mixed-effects model investigating the relationship between early-life rTL of *Rhinolophus ferrumequinum* juveniles and maternal age at conception (MAC).** Significant values ($p<0.05$) are highlighted in bold.

| Effects | Estimate | Std Error | t value | p-value |
|---|---|---|---|---|
| **Fixed Effects** | | | | |
| (Intercept) | -0.597 | 0.479 | -1.246 | 0.215 |
| **Age (in days)** | **-5.414** | **0.937** | **-5.778** | **<0.001** |
| **Age (in days)$^2$** | **2.687** | **0.981** | **2.739** | **0.006** |
| MAC | 0.002 | 0.010 | 0.210 | 0.834 |
| Sex (Male) | 0.055 | 0.115 | 0.480 | 0.632 |
| SMI | 0.040 | 0.035 | 1.142 | 0.254 |
| Sex (Male)*MAC | -0.004 | 0.014 | -0.297 | 0.766 |
| **Random Effects** | **Variance** | | | |
| qPCR Plate | 0.265 | | | |
| Individual identity | 0.068 | | | |
| Mother identity | 0.033 | | | |
| Year | 0.156 | | | |

**Table S14: Summary of mixed-effects model testing between and within mother age at conception (MAC) effects in *Rhinolophus ferrumequinum*.** Significant values ($p<0.05$) are highlighted in bold.

| Effects | Estimate | Std Error | t value | p-value |
|---|---|---|---|---|
| **Fixed Effects** | | | | |
| (Intercept) | -0.517 | 0.486 | -1.062 | 0.290 |
| **Age (in days)** | **-5.414** | **0.936** | **-5.783** | **<0.001** |
| **Age (in days)$^2$** | **2.649** | **0.980** | **2.703** | **0.007** |
| Mean MAC | -0.004 | 0.011 | -0.393 | 0.695 |
| Delta MAC | 0.099 | 0.055 | 1.809 | 0.071 |
| Sex (Male) | 0.008 | 0.116 | 0.068 | 0.946 |
| SMI | 0.037 | 0.035 | 1.045 | 0.296 |
| Sex (Male)*Mean MAC | 0.003 | 0.014 | 0.228 | 0.820 |
| **Sex (Male)*Delta MAC** | **-0.121** | **0.053** | **-2.275** | **0.024** |
| **Random Effects** | **Variance** | | | |
| qPCR Plate | 0.265 | | | |
| Individual identity | 0.059 | | | |
| Mother identity | 0.037 | | | |
| Year | 0.188 | | | |
| Residual | 0.478 | | | |

**Table S15: Fixed, random and variance-covariance matrices from a bivariate model of rTL and first year survival in female *Myotis myotis.*** Covariance and correlations were estimated at the yearly and among-individual levels. Estimates are given as the mode of the posterior distribution with 95% Highest Posterior Density credible intervals. For variance-covariance matrices, variances are shown on the diagonal, covariances in the bottom left and correlations in the top right. Significant covariances and correlations are shown in bold.

| **Response variables: rTL, Survival** | **Posterior mode** | **95% CI** | **Effective sample size** | **pMCMC** |
|---|---|---|---|---|
| *Fixed effects (n = 496)* | | | | |
| *rTL* | | | | |
| Intercept | -0.205 | [-2.048, 1.256] | 2651 | 0.705 |
| SMI | 0.021 | [-0.048, 0.083] | 3000 | 0.563 |
| *Survival* | | | | |
| Intercept | -1.929 | [-3.796, 1.145] | 3000 | 0.245 |
| SMI | 0.005 | [-0.043, 0.183] | 3000 | 0.299 |
| *Random effects* | | | | |
| qPCR plate | 0.280 | [0.110, 2.198] | | |
| *Variance-covariance matrices* | | | | |
| **Year sampled** | *rTL* | *Survival* | | |
| *rTL* | 0.314 [0.169, 1.260] | 0.461 [-0.188, 0.825] | | |
| *Survival* | 0.139 [-0.234, 0.941] | 0.486 [0.197, 2.104] | | |
| **Residual** | *rTL* | *Survival* | | |
| *rTL* | 0.577 [0.490, 0.700] | -0.017 [-0.188, 0.170] | | |
| *Survival* | -0.013 [-0.153, 0.124] | 1.000 [1.000, 1.000] | | |

**Table S16: Results of a bivariate Bayesian mixed-effects model jointly analysing juvenile relative telomere length (rTL) and first-year survival in female *Rhinolophus ferrumequinum*.** Estimates are given as the mode of the posterior distribution with 95% Highest Posterior Density credible intervals. rTL was modelled as a Gaussian trait with repeated measures, while survival to the following summer was modelled as a threshold (binary) trait measured once per individual. Random effects for qPCR plate and year were fitted for rTL only (*) to account for technical and temporal variation. Individual identity was specified with an unstructured (co)variance matrix, allowing estimation of among-individual variance in rTL, variance in individual rTL slopes, variance in survival, and their covariances. Correlation coefficients (r) are also shown. Significant covariances and correlations are shown in bold.

| **Response variables: rTL, Survival** | **Posterior mode** | **95% CI** | **Effective sample size** | **pMCMC** |
|---|---|---|---|---|
| *Fixed effects (n = samples, N = individuals)* | | | | |
| *rTL* | | | | |
| Intercept | -1.493 | [-3.027, 0.320] | 3000 | 0.116 |
| Age (centered) | -0.012 | [-0.027, 0.008] | 3000 | 0.323 |
| SMI | 0.105 | [-0.021, 0.222] | 3000 | 0.092 |
| *Survival* | | | | |
| Intercept | -0.208 | [-1.372, 0.695] | 3109 | 0.531 |
| SMI | 0.051 | [-0.022, 0.141] | 3084 | 0.175 |
| *Random effects* | | | | |
| qPCR plate* | 0.169 | [0.085, 0.349] | 3000 | |
| Year* | 0.297 | [0.092, 1.648] | 3200 | |
| Residual | 0.187 | [0.123, 0.294] | 2756 | |
| **Individual identity** | | | | |
| $V_{rTL}$ | 0.292 | [0.202, 0.402] | | |
| $V_{rTL:Slope}$ | 0.015 | [0.013, 0.018] | | |
| $V_{Surv}$ | 0.253 | [0.210, 0.308] | | |
| $COV_{rTL,rTL:Slope}$ | -0.001 | [-0.010, 0.009] | | |
| $COV_{rTL,Surv}$ | -0.006 | [-0.055, 0.046] | | |
| $COV_{rTL:Slope,Surv}$ | -0.001 | [-0.010, 0.008] | | |
| $r_{(rTL,rTL:Slope)}$ | -0.014 | [-0.148, 0.126] | | |
| $r_{(rTL,Surv)}$ | 0.011 | [-0.190, 0.167] | | |
| $r_{(rTL:Slope,Surv)}$ | -0.013 | [-0.149, 0.123] | | |

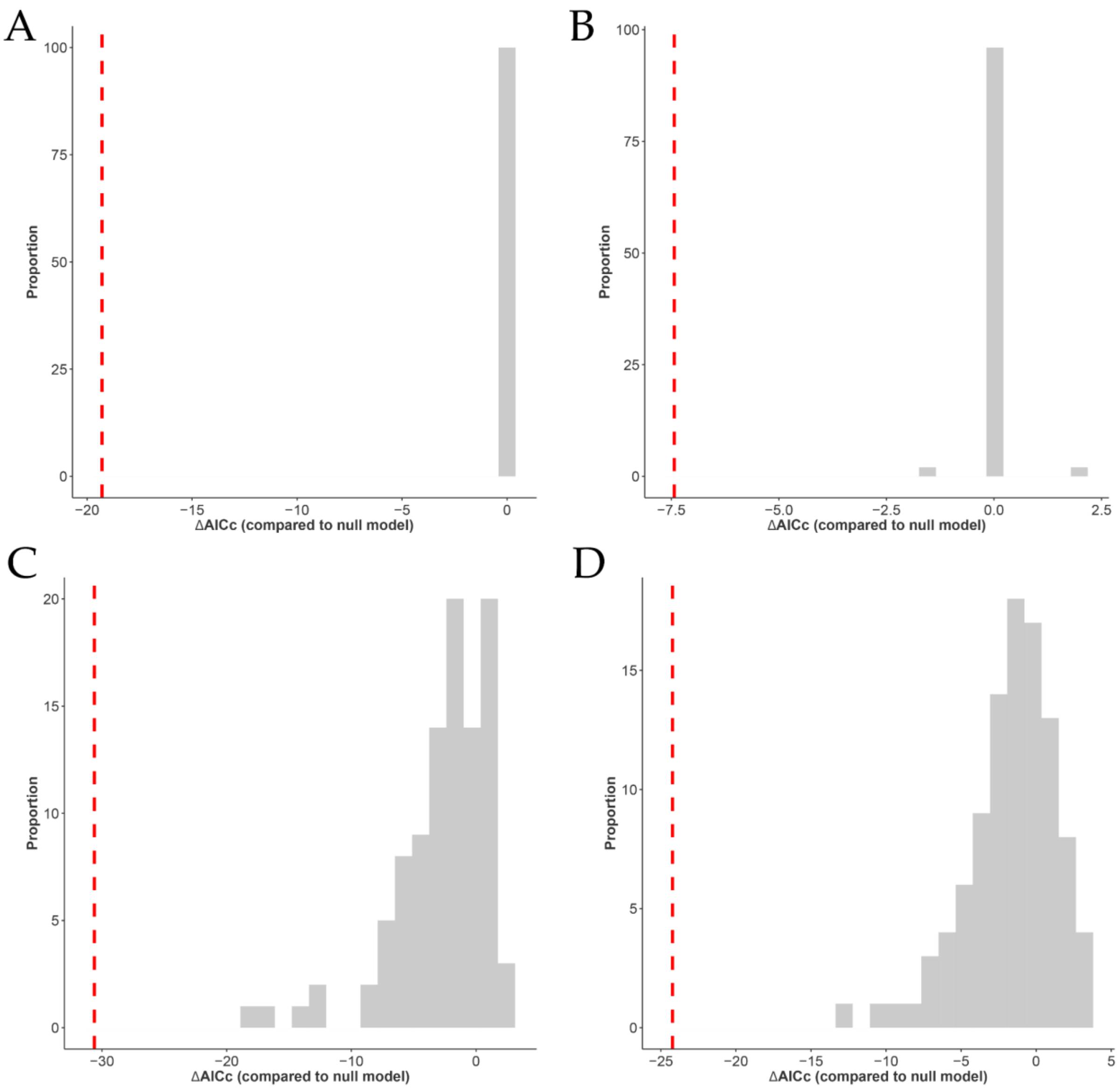


**Figure S1: Histograms representing the best AIC support for 100 randomised sliding window analyses for *Myotis myotis (*A) average daily rainfall (quadratic) and (B) average daily temperature (quadratic) analyses and *Rhinolophus ferrumequinum* (C) average daily rainfall (quadratic) and (D) average daily temperature (quadratic) analyses.** Red dashed lines represent the AIC support achieved with the real data. As some randomisation runs scored lower than the AIC support achieved with the real data, the average daily temperature (quadratic) pattern observed for *M. myotis* is considered a false positive.

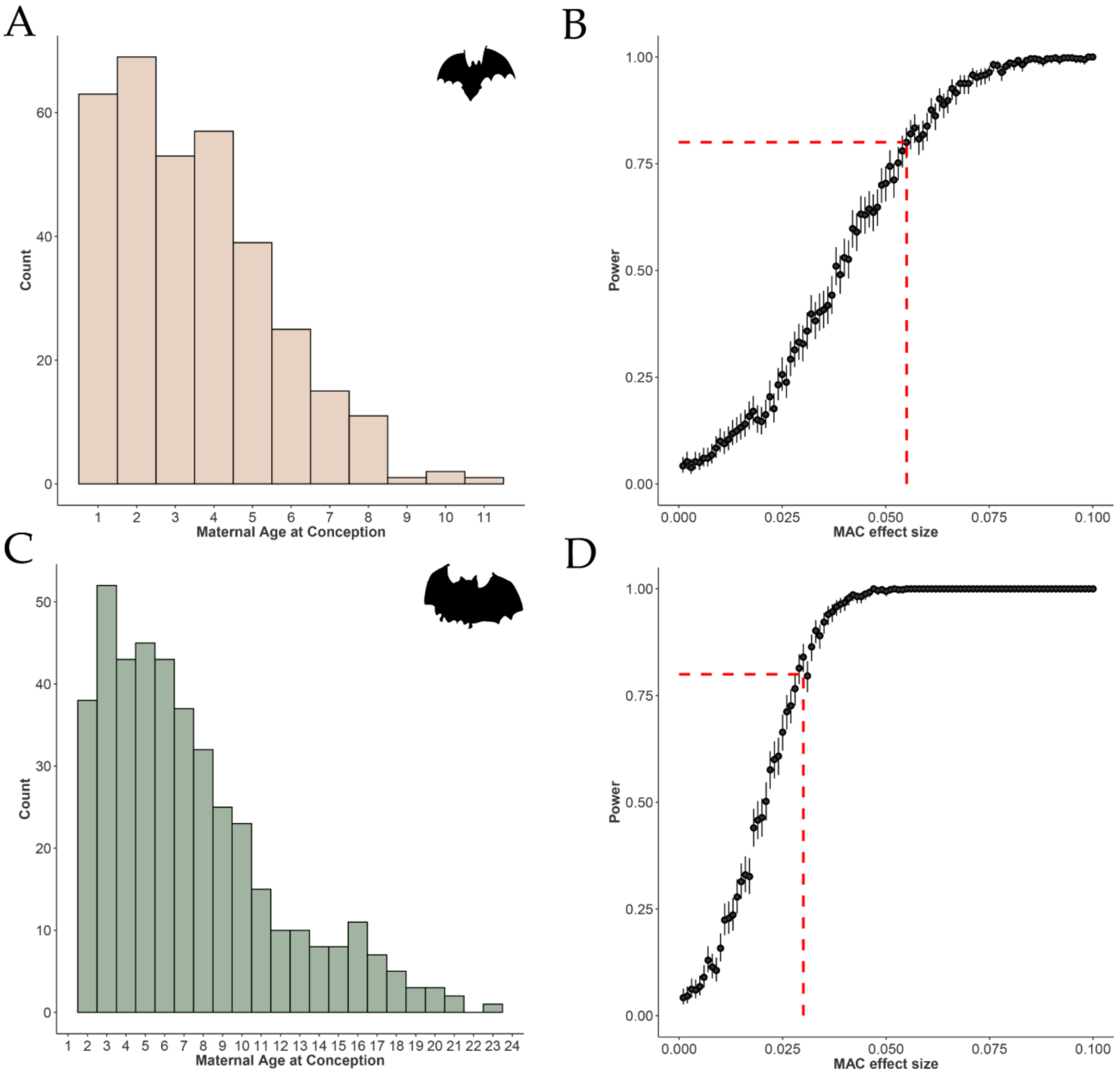


**Figure S2: Distribution of maternal ages and power analysis for detecting maternal age effects on juvenile *Myotis myotis* and *Rhinolophus ferrumequinum* telomere length.** (A) Histogram showing the distribution of maternal age at conception (MAC) for *M. myotis*. Maternal identity was assigned to 336 juveniles in this study, with maternal age estimated at the time of offspring conception. (B) Simulation-based power analysis for detecting MAC effects on juvenile relative telomere length in *M. myotis*. Power curves show the probability of detecting effects of increasing magnitude given the observed data and model structure. Red dashed lines indicate the effect size detectable with ≥80% power (MAC ≥ 0.05), corresponding to a correlation coefficient of r = 0.086. (C) Distribution of maternal age at conception for *R. ferrumequinum*. Maternal identity was assigned to 422 juveniles in this study. (D) Simulation-based power analysis for *R. ferrumequinum*, showing ≥80% power to detect MAC effect sizes ≥ 0.030, equivalent to a correlation coefficient of r = 0.131. Error bars represent 95% confidence intervals around simulated power estimates.
